# Genetic networks underlying natural variation in basal and induced activity levels in *Drosophila melanogaster*

**DOI:** 10.1101/444380

**Authors:** Louis P. Watanabe, Cameron Gordon, Mina Y. Momeni, Nicole C. Riddle

## Abstract

Exercise is recommended by health professionals across the globe as part of a healthy lifestyle to prevent and/or treat the consequences of obesity. While overall, the health benefits of exercise and an active lifestyle are well understood, very little is known about how genetics impacts an individual’s inclination for and response to exercise. To address this knowledge gap, we investigated the genetic architecture underlying natural variation in activity levels in the model system *Drosophila melanogaster*. Activity levels were assayed in the Drosophila Genetics Reference Panel 2 fly strains at baseline and in response to a gentle exercise treatment using the Rotational Exercise Quantification System. We found significant, sex-dependent variation in both activity measures and identified over 100 genes that contribute to basal and induced exercise activity levels. This gene set was enriched for genes with functions in the central nervous system and in neuromuscular junctions and included several candidate genes with known activity phenotypes such as flightlessness or uncoordinated movement. Interestingly, there were also several chromatin proteins among the candidate genes, two of which were validated and shown to impact activity levels. Thus, the study described here reveals the complex genetic architecture controlling basal and exercise-induced activity levels in *D. melanogaster* and provides a resource for exercise biologists.

## INTRODUCTION

Obesity is a disease associated with significantly higher all-cause mortality relative to normal weight (Carmienke *et al.* 2013; Flegal *et al.* 2013; Masters *et al.* 2013). This increase in mortality can be attributed largely to the elevated incidence of cardiovascular disease (Ebbert *et al.* 2014; Molica *et al.* 2015; Sisnowski *et al.* 2015), cancer (Berger 2014; Garg *et al.* 2014; Nakamura *et al.* 2014; Park *et al.* 2014), and diabetes (Nomura *et al.* 2010; Riobo Servan 2013; Polsky and Ellis 2015) observed in obese individuals. The increasing prevalence of obesity is a serious international public health issue that has warranted action from legislators worldwide (Hartemink *et al.* 2006). The upswing in rates of the disease throughout the United States caused the national medical expenditure dedicated to treating obesity-related illnesses in adults to increase by 29% between 2001 and 2015 (Biener *et al.* 2018). Therefore, strategies to counter the broadening obesity epidemic are needed to ensure the medical and financial wellbeing of society at large.

Exercise is among the most common treatments for obesity, which also include surgical procedures, medications, and other lifestyle modifications (Baretic 2013; Wyatt 2013; Kushner 2014; Martin *et al.* 2015). Given the relatively risk-free nature of exercise as a method of weight loss compared with many other treatment options, it is considered widely to be an essential component of treatment regimes for obesity (Mcqueen 2009; Laskowski 2012; Fonseca-Junior *et al.* 2013). In addition to treating obesity, exercise imparts a number of health benefits including improved muscle function (Andersen *et al.* 2015; Coen *et al.* 2015; Kim *et al.* 2015) and cartilage integrity (Tonevitsky *et al.* 2013; Breit *et al.* 2015; Blazek *et al.* 2016), increased insulin sensitivity (Mitrou *et al.* 2013; Brocklebank *et al.* 2015), and prevention of many chronic conditions (Booth *et al.* 2012). These benefits led exercise to be recognized as an important facet of a healthy lifestyle, with government agencies such as the U.S. Department of Health and Human Services issuing specific exercise recommendations for adults, youths, and children (e.g. adults should do at least 150 minutes of moderate-intensity, or 75 minutes of vigorous-intensity aerobic activity per week) (Dhhs 2008). Thus, exercise is an important component of many people’s lives, whether to treat obesity or to improve overall health.

Despite the growing relevance of exercise, there is a lack of clarity concerning a number of factors that influence its physiological effects (Karoly *et al.* 2012), most notably genetic background. Although exercise has gained considerable popularity as both a lifestyle choice and a treatment for obesity, there are large differences in how individuals respond to exercise, and it is not universally effective (Keith *et al.* 2006; Mcallister *et al.* 2009). In fact, exercise provides no metabolic improvements to certain individuals (Bouchard *et al.* 1999; Bouchard *et al.* 2011; Stephens and Sparks 2015), revealing an extreme disparity in exercise response that can likely be accounted for, at least in part, by genetic variation. Moreover, existing data suggest a relationship between exercise-induced improvements to muscle metabolism and exercise performance in humans (Larew *et al.* 2003). Thus, while some genes have been identified as contributors to physical activity traits of an individual (Stubbe *et al.* 2006; De Geus *et al.* 2014), the genetic architecture controlling exercise responses has yet to be characterized.

As an emerging model organism for exercise studies, *Drosophila melanogaster* possesses several characteristics advantageous for elucidating the relevant genetic architecture (Piazza *et al.* 2009; Tinkerhess *et al.* 2012; Sujkowski *et al.* 2015; Mendez *et al.* 2016; Watanabe and Riddle 2017). Traditional obstacles for exercise studies, including difficulties in controlling for essential variables such as age, sex, fitness, and diet, can be addressed using *D. melanogaster*. Furthermore, Drosophila is an established model system with a well-characterized genome and ample tools for genetic studies. These tools include the Drosophila Genetics Reference Panel 2 (DGRP2), a fully sequenced population of 200 genetically diverse isogenic lines for quantitative genetic studies (Mackay *et al.* 2012; Huang *et al.* 2014), in addition to large collections of mutants and RNAi knockdown lines. Furthermore, a high degree of genetic (Schneider 2000) and functional (Hewitt and Whitworth 2017) conservation exists between Drosophila and humans, particularly in areas such as energy-related pathways (Edison *et al.* 2016) and disease genes (Pandey and Nichols 2011; Ugur *et al.* 2016), which often allows findings from Drosophila to be translated to mammalian model systems. Together, these features make Drosophila an excellent choice for studies of exercise genetics.

Several innovative studies demonstrate that exercise treatments of Drosophila produce significant physiological and behavioral responses, including increased lifespan and improved climbing ability (Piazza *et al.* 2009; Tinkerhess *et al.* 2012; Sujkowski *et al.* 2015; Mendez *et al.* 2016; Watanabe and Riddle 2017). For example, treatments with the Power Tower, which prompts exercise by exploiting the negative geotaxis of Drosophila by repeatedly dropping their enclosures, causing the animals to fall to the base and attempt another climb, improves mobility in aging animals (Piazza *et al.* 2009). The TreadWheel exploits negative geotaxis through slow rotation of fly enclosures to stimulate a response; responses to prolonged exercise on the TreadWheel include, for example, changes in triglyceride and glycogen levels in the animals (Mendez *et al.* 2016). The Rotating Exercise Quantification System (REQS) is an offshoot of the TreadWheel, which is able to record the activity levels of flies as they exercise, facilitating the comparison of different exercise regimes and allowing for the normalization of exercise levels (Watanabe and Riddle 2017). The REQS validation study also demonstrated that there is significant variability among different Drosophila genotypes in how they respond to the rotational exercise stimulation (Watanabe and Riddle 2017), suggesting that genetic factors contribute to the difference in exercise levels observed.

In this study, we investigate the genetic factors contributing to the level of exercise induced through rotational stimulation. Using the REQS, we measured basal activity levels (without rotation) as well as induced exercise levels (with rotation) in 161 genetically diverse strains from the DGRP2. Next, we used a genome-wide association study (GWAS) to identify the genetic variants responsible for the approximately 10-fold variation in activity levels observed within the DGRP2 lines. We identified over 100 annotated genes that contribute to basal and induced activity levels. The loci that control activity levels are different for the untreated and exercise-treated conditions and often also differ between males and females. Additional characterization of candidate genes validate the results of our GWAS and confirm that genes with functions in the central nervous system as well as some chromatin proteins impact Drosophila activity levels. Together, our findings provide key insights into the number and types of genetic factors that control basal and exercise-induced activity levels, provide an array of candidate genes for follow-up studies, and identify chromatin modifiers as a new class of proteins linked to exercise.

## MATERIALS AND METHODS

### Drosophila Lines and Husbandry

The DGRP2 fly lines used in this study were obtained from the Bloomington Drosophila Stock Center or from our collaborator Dr. Laura Reed (University of Alabama). Fly lines for the follow-up analysis of candidate genes (Supplemental Table S1) were obtained from the Bloomington Drosophila Stock Center. Drosophila were grown on media consisting of a cornmeal-molasses base with the addition of Tegosept, propionic acid, agar, yeast, and Drosophila culture netting (Mendez *et al.* 2016). All flies used for the assays were reared in an incubator at 25°C and ∼70% humidity with a 12-hour light-dark cycle for a minimum of three generations prior to the start of the experiments. To minimize density effects, animals were grown in vials established by mating seven male to ten female flies for each line. The resulting progeny were collected as virgins, aged 3-7 days, separated by sex, and then used for the exercise experiments.

### Exercise Quantification Assay

Basal and induced exercise activity levels were determined for a total of 161 strains (155 basal and 151 induced) from the DGRP2 using the REQS (Mackay *et al.* 2012; Huang *et al.* 2014; Watanabe and Riddle 2017; Watanabe and Riddle 2018) (Supplemental Table S2). For each DGRP2 line, 100 male and 100 female virgin flies aged 3-7 days were used. The animals were anesthetized with CO_2_, divided into groups of 10 (n=10 groups per sex per line), and loaded into the vials of the REQS at 9AM [for additional details see (Watanabe and Riddle 2018)]. The REQS was moved into the incubator at 10AM, and the vials were rotated to a vertical orientation similar to how flies are reared in a laboratory setting. The animals were allowed to recover from anesthetization for one hour. The basal activity level of the animals was measured from 11AM to 12PM, keeping the REQS in a static position without rotation. At 12PM, the REQS’ rotational feature was turned on at 4 rotations per minute (rpm) to induce exercise in the animals, and activity levels were monitored during this exercise until 2PM. Measurements were taken at five-minute intervals during both the basal and induced activity phases.

### Statistics

Basal and induced activity levels were calculated as the average activity level per five-minute interval of each vial/genotype/sex combination. A GLM (general linear model with gamma log-link) to investigate the impact of the factors vial, genotype, sex, and treatment on activity levels was performed using SAS9.4 software (Inc 2013). As the initial analysis showed no effect of “vial”, “vial” was removed from the final model. Descriptive statistics were generated in SPSS25 (Ibm 2017) and R (R Development Core Team 2018). Custom Perl scripts were used for the SNP classification analysis in addition to R (R Development Core Team 2018).

### Genome Wide Association Study (GWAS)

Basal and induced activity levels were separately analyzed by calculating the average activity level per five-minute interval of each genotype/sex combination. These phenotypic values (Supplemental Table S3) were used for two separate GWASs using the DGRP2 webtool (http://dgrp2.gnets.ncsu.edu/) developed by Dr. Trudy Mackay (Mackay *et al.* 2012; Huang *et al.* 2014). Genetic variants that met a significance threshold of p<10^-5^ in any of the analyses (mixed model, simple regression model, female data, male data, combined data, and sex difference analysis) were considered candidate loci (Supplemental Table S4). Candidate genes for follow-up and validation were selected based on significance level, mutant availability from Drosophila stock centers, and reports on FlyBase (Gramates *et al.* 2017) consistent with phenotypes that might be linked to exercise/activity.

### Gene Ontology (GO)

GO analysis was performed on the genes associated with the genetic variants that met the significance threshold of p<10^-5^ separately for both the basal and induced GWAS results. Genetic variants lacking association with a specific gene were removed from the list, and duplicate genes were removed as well. FlyBase gene IDs were retrieved for the gene sets using The Database for Annotation, Visualization and Integrated Discovery (DAVID) ID conversion function (Huang Da *et al.* 2009b; Huang Da *et al.* 2009a). GO analysis was carried out using PANTHER (Protein Analysis Through Evolutionary Relationships) (Mi *et al.* 2013; Mi *et al.* 2017). The “biological process” and “cellular component” enrichment terms were used.

### Functional analysis of candidate genes

Candidate genes were selected for follow-up experiments based on their *p*-values in the GWASs, as well as based on the annotation available on FlyBase. For knockdown of *Su(z)2* and *Jarid2*, UAS-controlled RNAi constructs (*y^1^ sc* v^1^ sev^21^; P{y*[*+t7.7*] *v*[*+t1.8*]*=TRiP.HMS00281}attP2/TM3, Sb^1^* for *Su(z)2* [Bloomington stock 33403]; *y^1^ sc* v^1^; P{y*[*+t7.7*] *v*[*+t1.8*]*=TRiP.HMS00679}attP2* for *Jarid2* [Bloomington stock 32891]) were combined with a muscle-specific or central nervous system specific driver (muscle: *P{w*[*+mC*]*=UAS-Dcr-2.D}1, w^1118^; P{w*[*+mC*]*=GAL4-Mef2.R}R1* [Bloomington stock 25756]; central nervous system: *P{w*[*+mW.hs*]*=GawB}elav*[*C155*] *w^1118^; P{w*[*+mC*]*=UAS-Dcr-2.D}2* [Bloomington stock 25750]) (Perkins *et al.* 2015). Offspring from these crosses carrying the UAS construct as well as the GAL4 driver were collected and aged 3-5 days prior to the start of the experiment. Basal and exercise-induced activity levels were measured as described above, with the following modification: Activity measures were collected both in the morning (10AM-1PM) and afternoon (4PM-7PM), with time used as a blocking factor.

### Data availability

All data necessary for confirming the conclusions of this article are represented fully within the article and its tables, figures, and supplemental files, with the exception of the genotype data for the DGRP2 population and the GWAS model. This information can be found at http://dgrp2.gnets.ncsu.edu/.

## RESULTS

### The DGRP2 shows extensive variation in basal and exercise-induced activity levels

To investigate the genetic basis of variation in activity levels, both basal and exercise induced, we focused on the DGRP2 collection of Drosophila strains. The DGRP2 is a collection of 200 inbred lines of Drosophila derived from wild-caught females, representing genetic variation that is present in a natural population. To measure activity levels, we used the REQS, as it allowed us to record basal activity of the animals without rotation and induced activity levels during rotation. The rotation induces exercise (higher activity levels) through the animals’ negative geotaxis response. Using 161 strains from the DGRP2, we measured basal activity of the animals as well as the activity during rotationally-induced exercise in a single experiment (Figure 1). The output from the REQS is the average activity level per 10 flies per five-minute interval, which was estimated based on a one-hour recording for the basal activity and a two-hour recording for the exercise activity.

**Figure 1.**
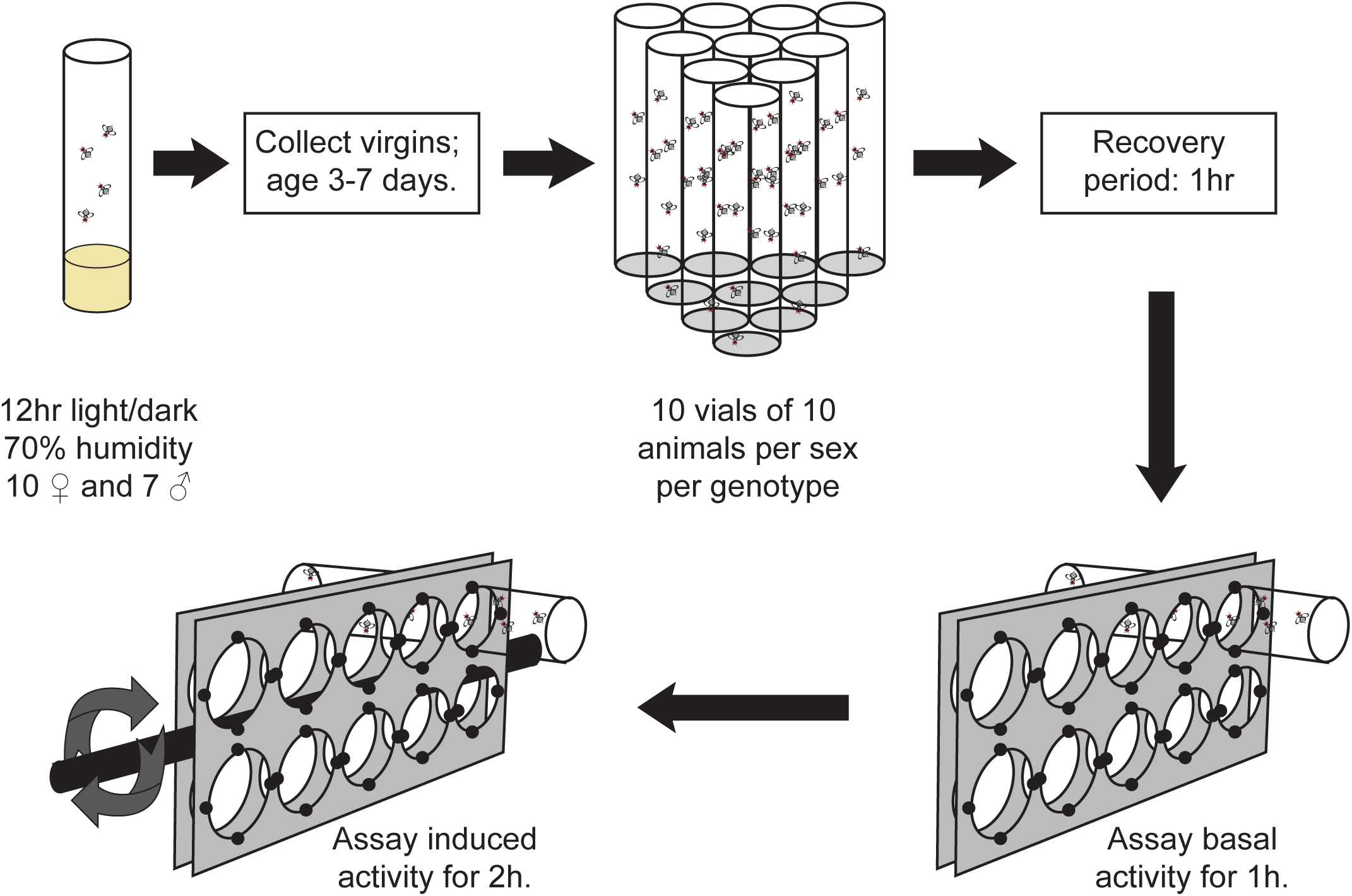
Experimental setup. Diagram illustrating the experimental setup, how animals were processed, and data were collected.

Figure 2 shows the results from this set of experiments, with data from females (A+B) and males (C+D) shown separately (see also Figure S1). All activity measures described are given as crossings/five minutes/fly. We found a wide range of average activity levels for both basal and exercise-induced activity. The highest level of basal activity in males was found in line 57, which exhibited 144.5 +/- 9.4 activity units (mean +/- SEM), while for females line 595 had the highest basal activity with 58.55 activity units (+/- 3.9). The highest performing line was the same for exercise-induced activity in both sexes, line 808 with an average of 155.25 activity units (+/- 3.56) in males and an average of 133.94 activity units (+/- 3.60) in females. Interestingly, for males the lowest basal and induced activity levels were measured both in line 383 with 0.275 activity units (+/- 0.1) for basal activity and 1.63 activity units (+/- 0.79) for exercise-induced activity. The low performer in the females was different between basal and induced activity: Line 390 females had the lowest basal activity with 0.53 activity units (+/- 0.14), while line 32 with an average of 1.22 activity units (+/- 0.26) had the lowest exercise-induced activity level. Looking across all factors, the variation in mean activity levels illustrated in Figure 2 range from a low of 0.275 +/- 0.1 activity units (line 383, basal males) to a high of 155.25 +/- 3.56 activity units (line 808, induced males). Thus, our highest activity measurement showed an approximately 500-fold increase from the lowest measurement, demonstrating that there is extensive variation in animal activity based on genotype, sex, and treatment within the DGRP2 population.

**Figure 2.**
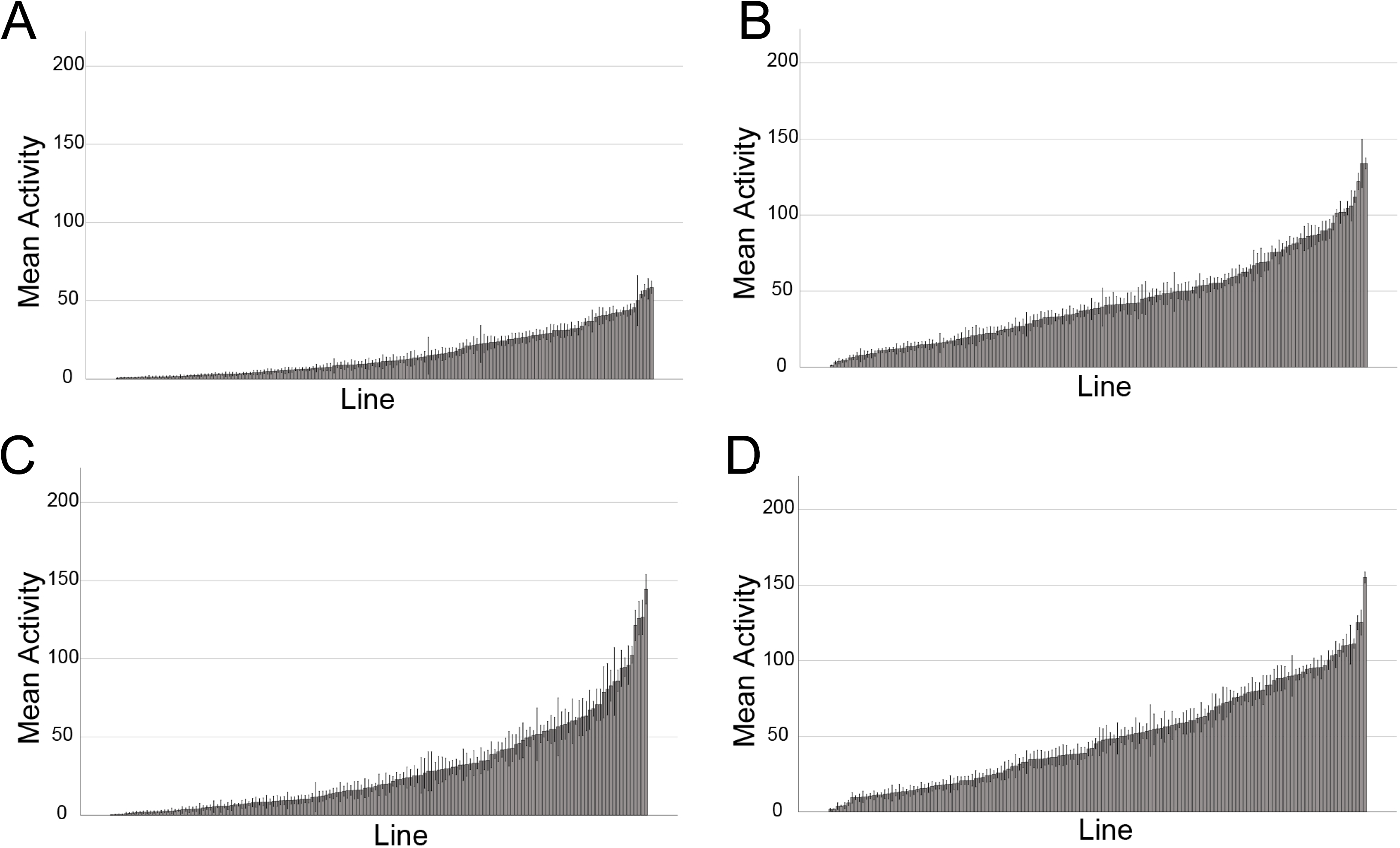
Activity levels within the DGRP2 population vary significantly. This set of bar graphs illustrate the variability in activity levels among the different lines of the DGRP2 population. In each graph, the lines are ordered based on their activity levels from smallest to largest (X-axis), with mean activity levels (average number of beam crossings recorded by the activity monitor in a 5-minute interval) on the Y-axis. The error bars are SEM. **A.** Bar graph showing line means for basal activity – female. **B.** Bar graph showing line means for induced activity – female. **C.** Bar graph showing line means for basal activity – male. **D.** Bar graph showing line means for induced activity – male.

Several general trends can be discerned from the bar graphs in Figure 2: 1) Across the 161 lines, female flies tend to have a lower basal activity level than males (compare A to C); 2) rotational stimulation tends to increase activity levels (compare A to B and C to D), and 3) while basal activity levels tend to be lower in females, exercise activity levels tend to be more similar between the sexes (compare A and C to B and D). However, there are exceptions to these general trends. For example when comparing male to female activity, of the 155 basal lines measured, 27 lines displayed significantly higher activity levels in males, but there is one genotype (line 83) which exhibited higher activity levels in females (40.35 female activity, 4.87 male activity, p=0.044). Similarly, for the induced activity levels, 12 lines showed significantly greater activity in males than in females, while there was one line that showed the opposite trend, higher activity levels in females than in males (line 796, 122.01 female activity, 75.6 male activity, p<0.001). When observing the effects of exercise, as expected 41 genotypes demonstrated significantly increased activity with only three lines showing decreased activity. Thus, most genotypes showed increased activity when rotated, and males exhibited higher activity levels than females.

### Exercise-induced activity measures are strongly correlated between males and females of the same genotype

While the graphs presented in Figure 2 provide an assessment of the variability in activity phenotypes among the DGRP2 strains, they do not reveal how activity levels between males and females of the same strain relate to each other, nor do they reveal the relationship between basal and exercise-induced activity levels within the same strain/sex. To address these questions, we used scatter plots and determined the correlations of activity measures between males and females of the same strain, as well as the correlations between basal and exercise-induced activity separately for both sexes (Figure 3). We find that the Pearson correlation coefficient between male and female measures for basal activity levels is 0.32 (p<0.01), indicating a weak positive relationship (Figure 3A). The correlation between male and female measures for induced exercise is 0.831 (p<0.01), suggesting a strong relationship between the two measures (Figure 3B). When we examine the relationship between basal and exercise-induced activity levels, we find that in females, there is a moderately strong positive relationship between the two measures (Pearson correlation coefficient 0.512, p<0.01; Figure 3C). In males, the correlation between basal and exercised-induced activity is somewhat weaker, with a Pearson correlation coefficient of 0.350 (p<0.01; Figure 3D). These results suggest that all measures show some degree of positive correlation. The strongest correlation is seen among both sexes for the exercise-induced activity phenotype, possibly reflecting the fact that this measure is most strongly impacted by the animals’ overall physical abilities.

**Figure 3.**
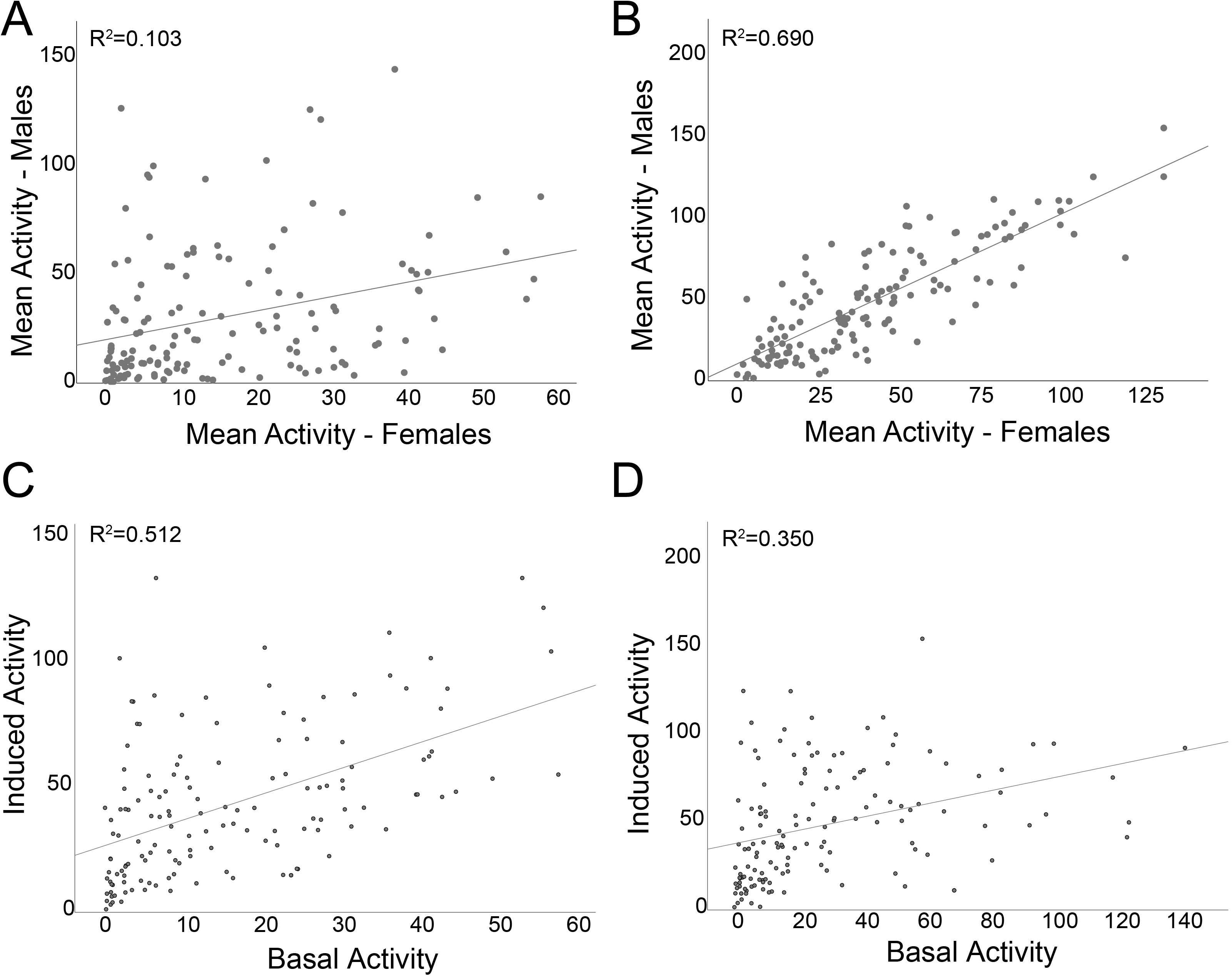
Exercise-induced activity measures are strongly correlated between males and females of the same genotype. This set of scatter plots examines the relationship between activity measures of males and females of the same genotype (A – basal activity; B – exercise-induced activity) and between the two activity measures in either females (C) or males (D). Mean activity levels are plotted as average number of beam crossings recorded by the activity monitor in a 5-minute interval. A linear regression line is shown in all plots, along with the correlation coefficient. **A.** Scatter plot showing the relationship between basal activity levels in females (X-axis) and in males (Y-axis). **B.** Scatter plot showing the relationship between induced activity levels in females (X-axis) and in males (Y-axis). **C.** Scatter plot showing the relationship between basal (X-axis) and exercise-induced (Y-axis) activity levels in females. **D.** Scatter plot showing the relationship between basal (X-axis) and exercise-induced (Y-axis) activity levels in males.

Because the DGRP2 strains have been used to investigate a wide range of phenotypes by the Drosophila research community, we were able to also look at the connection between animal activity levels and lifespan. Generally, it is thought that a more active lifestyle and overall higher activity levels lead to a healthier and longer life (Reimers *et al.* 2012). Surprisingly, in the DGRP2 strains, we find no correlation between the basal activity levels of the animals measured by the REQS and the lifespan reported by Durham and colleagues [Pearson’s correlation: 0.029; p= 0.7417; Figure S2A; (Durham *et al.* 2014)]. Similarly, if we examine the relationship between exercise-induced activity levels and lifespan, there is no significant correlation (Pearson’s correlation: −0.136; p=0.1245; Figure S2B). This lack of relationship between activity levels and lifespan is unexpected and deserves further investigation.

### Sex, genotype, and exercise treatment impact activity

Next, we investigated the factors influencing the variation in activity levels we observed in Figure 2 utilizing a general linear model (GLM) analysis. Specifically, the GLM examined the impact of treatment (with or without rotation), sex, genotype, as well as the interactions between these factors. The results indicate that activity levels were significantly impacted by treatment, illustrating that rotation indeed is able to increase activity levels above baseline in this genetically diverse population of fly strains (p<0.001, Table 1). In addition, sex and genotype significantly impacted activity levels, as suggested by the descriptive data illustrated in Figures 2 and S1 (p<0.001). Interaction effects between treatment, sex, and genotype also impacted the activity levels measured by the REQS, indicating that to be able to predict activity phenotypes, sex, treatment, and genotype must all be considered together (Table 1). The descriptive statistics combined with the GLM analysis thus indicate that for the activity phenotype there is tremendous variation between the DGRP2 lines, some of which is due to genetics (genotype effect). This finding suggested that individual genes underlying the variation in activity levels might be identified by a GWAS.

**Table 1.** Treatment, sex, and genotype show strong effects on activity levels. General linear model analysis of variance reveals the impact of treatment (+/- rotational stimulation), sex, and genotype on the activity phenotype measured.

| Source | Type III |  |  |
| --- | --- | --- | --- |
|  | Wald Chi-Square | df | p-value |
| (Intercept) | 115810.796 | 1 | 0.000 |
| Genotype | 8550.945 | 160 | 0.000 |
| Sex | 377.098 | 1 | 0.000 |
| Treatment | 3273.344 | 1 | 0.000 |
| Genotype * Sex | 2342.484 | 155 | 0.000 |
| Sex * Treatment | 109.291 | 1 | 0.000 |
| Genotype * Treatment | 2367.982 | 145 | 0.000 |
| Genotype * Sex * Treatment | 857.944 | 140 | 0.000 |
| Dependent Variable: Activity |  |  |  |
| Model: (Intercept), Genotype, Sex, Treatment, Genotype * Sex * Treatment, Genotype * Sex, Sex * Treatment, Genotype * Treatment |  |  |  |

df: degrees of freedom

### GWAS identifies over 400 genetic variants impacting activity levels

To identify the genetic factors impacting basal and exercise-induced activity levels in the DGRP2 population, we carried out a GWAS. The analysis was run separately for the basal and exercise-induced activity levels (155 and 151 lines respectively) using the DGRP2 GWAS Webtool (http://dgrp2.gnets.ncsu.edu/). The webtool uses two different models for the analysis: a simple regression model and a more complex mixed model. The analysis is carried out for male data only, female data only, combined data from both sexes, and for the difference between sexes. Together, this set of GWASs identified over 400 genetic variants [single nucleotide polymorphisms (SNPs), multiple nucleotide polymorphisms (MNPs), and deletions (DEL)] that impact basal and exercise-induced activity levels (p<0.005; Figures 4 and S3, Tables 2 and S4), illustrating the importance of genetic factors for exercise phenotypes.

**Figure 4.**
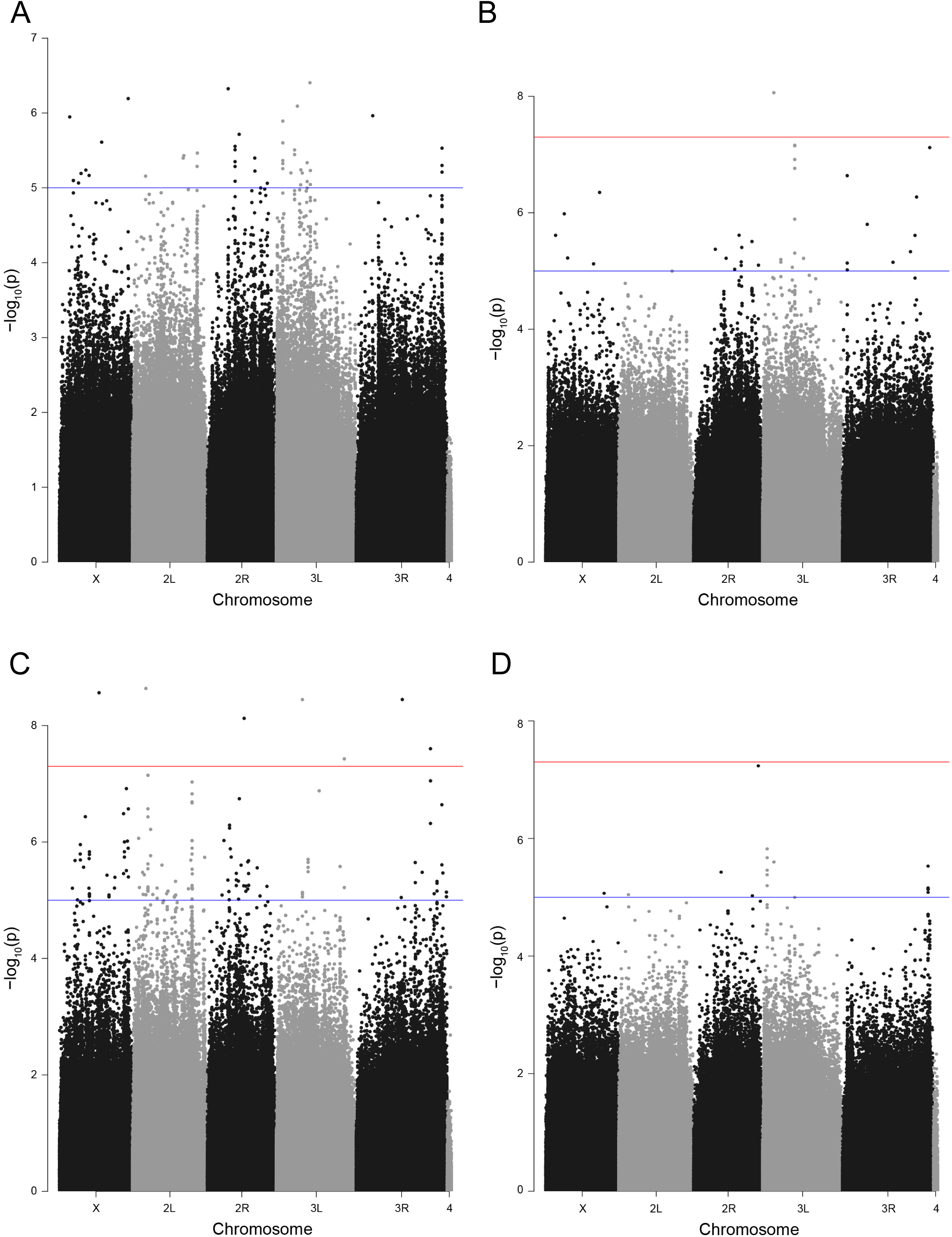
GWAS identifies SNPs associated with basal and induced activity levels. Chromosomal location of SNPs (X-axis) are plotted against the negative log of the p-value testing for the likelihood of the SNP being associated with the measured phenotype (mixed model; Y-axis). The blue line in each plot marks the p=10^-5^ significance level, while the red line marks p=10^-7^. **A.** Manhattan plot for basal activity – female. **B.** Manhattan plot for induced activity – female. **C.** Manhattan plot for basal activity – male. **D.** Manhattan plot of induced activity – male.

**Table 2.**
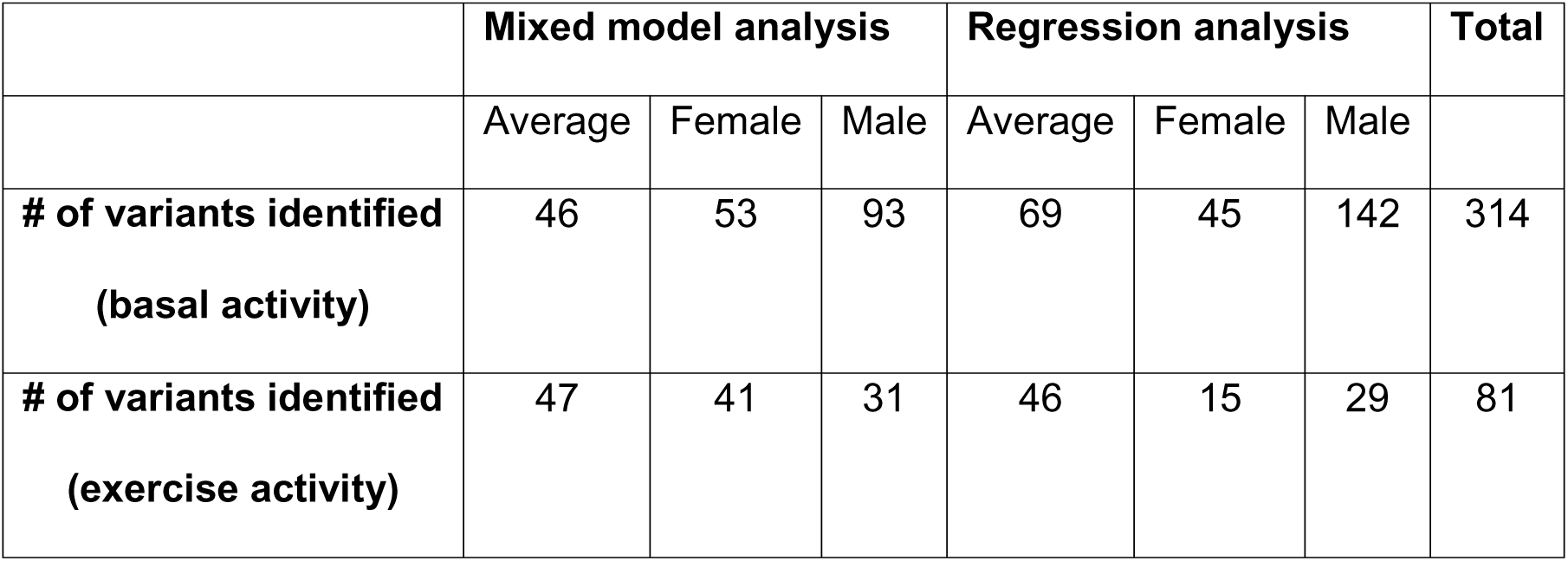
Summary of GWAS results. The table lists the number of genetic variants detected as significant by the different analysis types used in this study.

The genomic variants identified by the GWAS are distributed throughout the genome, and with the exception of the small 4^th^ chromosome, all chromosome arms contain genetic variants contributing to the basal and induced exercise activity phenotypes (Table S4). Comparing the chromosomal distribution of the genetic variants identified as significant in the basal activity or induced exercise activity analysis to that of the overall distribution of variants using chi-square analysis, we found no significant deviations from the expectation, indicating that the variants are not clustered in any particular way in the genome. However, there are small areas of linkage disequilibrium, where significant variants are clustered on chromosomes 2L and 3L (basal analysis) and chromosomes 2R, 3R, and 3L (induced analysis; Figure S4). Overall, the results of this GWAS demonstrate that a large number of loci contribute to the two activity phenotypes measured and that the genetic architecture underlying the phenotypes is complex.

### Basal activity variant distributions differ significantly from genome-wide set

Next, we investigated what types of genetic variants were identified as significant in the GWAS. In order to compare the classifications of the variants from the basal and induced analyses to the entire genome, the variants from the induced and basal activity were first characterized based on their genomic context (introns, exons, upstream, downstream, UTR, and unknown). We then compared the classification distributions from the basal and induced analyses to a genome-wide set using a chi-square test. We found that only the basal activity variant distributions were significantly different from the genome-wide set (p=5.413*10^-7^) with a greater number of exons, upstream, and unknown elements (Figure 5). The classification for the variants contributing to induced exercise activity were not significantly different from the genome-wide classification of variants, possibly due to the smaller number of variants detected in this analysis and a concomitant reduced power to detect differences. Thus, the GWASs carried out here identify a diverse set of genetic variants as contributing to the activity phenotypes under study, and the overrepresentation of exon variants in the basal activity analysis suggests that these variants might indeed present genes important for animal activity and exercise.

**Figure 5.**
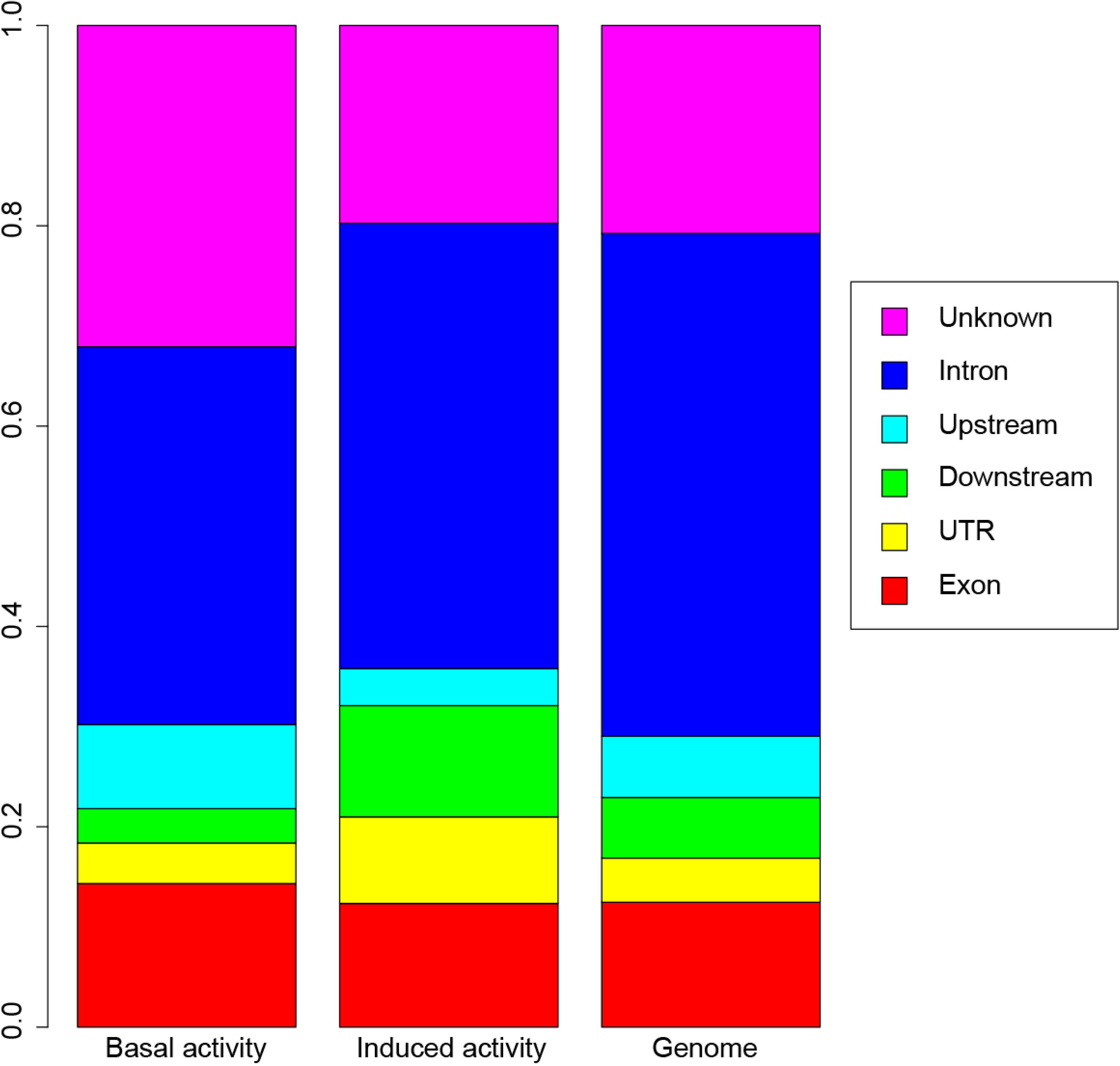
The genetic variants identified by the GWAS for basal activity are biased towards exons and upstream genetic elements. The genetic variants identified as significantly associated with either the basal (left) or exercise-induced (middle) activity were classified based on their sequence context and compared to the genome-wide set of variants forming the basis of this GWAS (right). The set of variants identified in the basal activity GWAS is significantly different from the genome-wide set (p=5.42*10^-7^; Chi-square test).

### Genetic variants impacting activity levels differ between males and females

The genetic variants identified as significantly associated with the two activity phenotypes differed between the analyses, some linked to activity in the males, some linked to activity in females, and some variants linked to the difference in activity levels between males and females. While the separate analyses of both male and female data lead to the identification of genetic variants, more genetic variants were identified in males than in females, and the combined analysis recovered more variants identified in males than in females. For example, the mixed model analysis of basal activity levels identified 53 genetic variants in females, 93 genetic variants in the males, and in the combined analysis, 46 variants are identified as significant, 22 of which overlap with the variants identified using the male data, while only one variant also occurs in the female analysis (p<5*10^-5^). These results illustrate the importance of collecting data in both sexes, as the genetic factors contributing to both basal and induced exercise activity levels differ between males and females. The results also suggest that the difference between males and females is under genetic control.

### GWAS discovers positive and negative effect variants impacting activity levels

The genetic variants identified as significantly associated with basal and/or induced exercise activity include loci with positive and negative effects on the phenotype. In the analysis of the induced exercise activity data, only genetic variants with negative impacts on activity levels were identified in either sex. Examining the data for basal activity, in males, the majority of variants negatively impacted basal activity, and only 1.5% of variants (three out of 195; mixed model) increased activity levels. These rare genetic variants leading to increased activity levels were associated with the genes *bdg* and *slo*. In females, the results are similar, with the majority of variants (63; mixed model) impacting basal activity levels negatively and positive effect variants being rare (three variants associated with *CG32521* and *CG8420*). Overall we found that the vast majority of our discovered variants resulted in a negative effect on activity.

### Terms related to the central nervous system and muscle function are over-represented in the GO analysis

Next, we focused on the genetic variants identified in the GWAS as significant that were associated with genes and asked what types of genes contributed to the basal and induced activity phenotypes (146 genes for basal activity; 47 genes for induced exercise activity). To do so, we explored the gene ontology (GO) terms associated with the gene sets linked to basal and induced activity levels. In order to determine which gene classes were over- or under-represented, GO analysis was carried out using the PANTHER (Protein Analysis Through Evolutionary Relationships) tools (Mi *et al.* 2013; Mi *et al.* 2017). Specifically, enrichment analysis was used to identify biological processes, cellular components, or molecular functions over-represented within the GWAS gene set relative to the genome as a whole.

For the GO term enrichment analysis for the genes contributing to basal activity levels, 30 GO terms were identified in the “biological process” category and 17 in the “cellular component” category. Among the “biological process” and “cellular component” GO terms, the largest fold enrichments were seen for two terms related to neurons, “axonal growth cone” (26.24-fold enrichment; p=3.3*10^-4^) and “neuron recognition” (7.51-fold enrichment, p=5.79*10^-5^) (Figure 6). Other significantly enriched GO terms include “neuromuscular junction”, “synaptic transmission”, and “behavior” (Figure 6). The GO term enrichment analysis of the gene set associated with exercise-induced activity identified a 27-fold enrichment for genes involved in the Alzheimer disease-presenilin pathway (p=2.01*10^-4^). Thus, the GO term enrichment analysis identifies a variety of terms associated with neuronal function and behavior as characterizing the gene set involved in controlling basal and exercise-induced activity levels.

**Figure 6.**
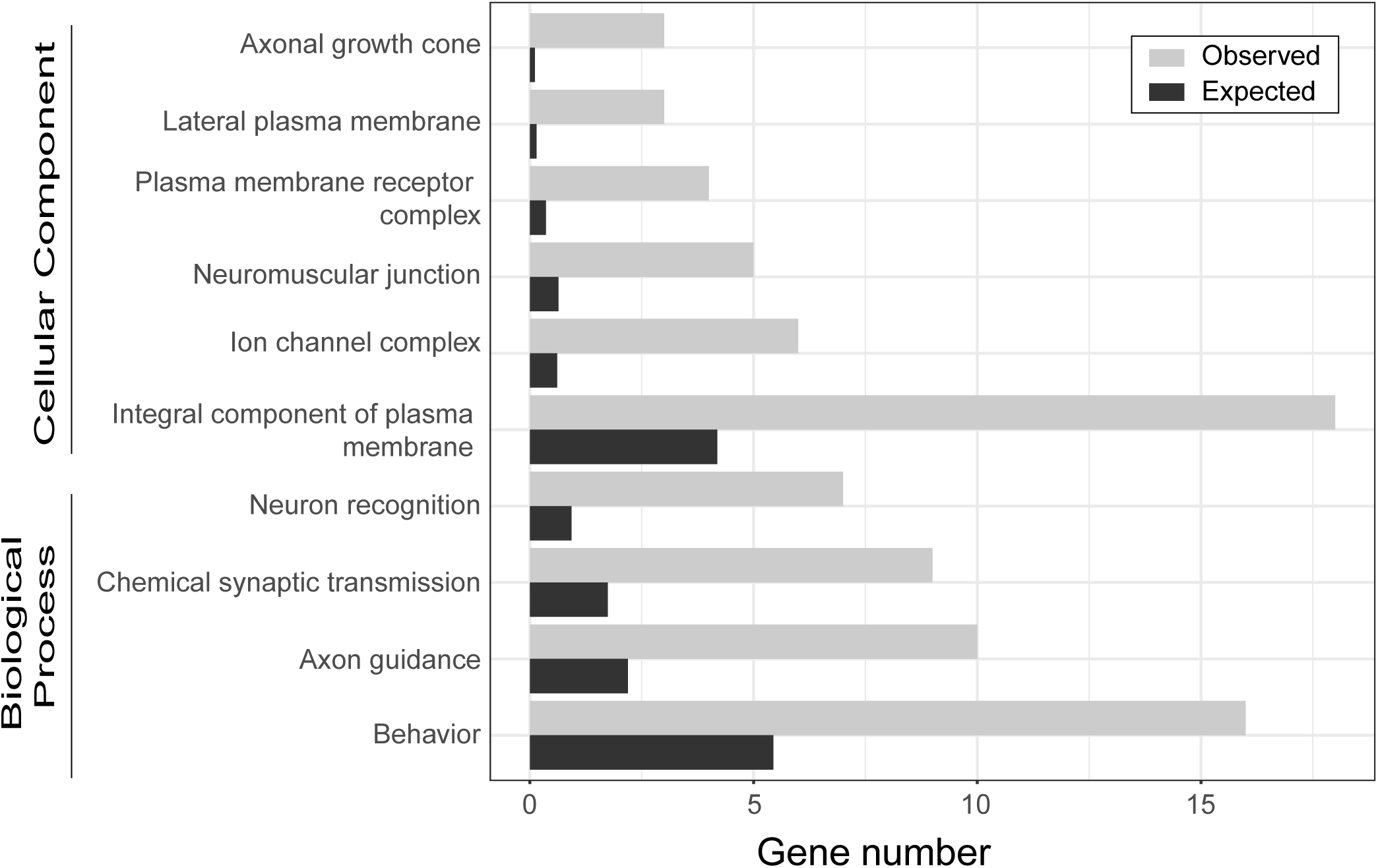
GO analysis highlights the importance of axonal growth cone and neuron recognition for activity levels. Genetic variants identified in the GWAS for basal analysis were subjected to GO enrichment analysis. Only high-level GO terms meeting the significance threshold are shown (for complete results and individual p-values, see Supplemental Table S5).

### Candidate gene analyses support the GWAS results and suggest that the chromatin proteins SU(Z)2 and JARID2 impact animal activity

In order to assess the success of the GWAS analysis, we surveyed the information available on FlyBase (Gramates *et al.* 2017) to determine if altered activity phenotypes have been described associated with the candidate genes identified here. Mutants in 25 of the candidate genes have been described as either “flightless”, or “flight defective” on FlyBase (Gramates *et al.* 2017), five were described as “uncoordinated,” and one (*Cirl*) was described as “hyperactive” and “sleep defective” (Van Der Voet *et al.* 2015). In our follow-up studies, we focus on two proteins involved in *polycomb* mediated gene regulation, *Su(z)2*, a member of the Polycomb Repressive Complex 1 [PRC1] (Wu and Howe 1995; Lo *et al.* 2009; Nguyen *et al.* 2017), and *Jarid2*, a Jumonji C domain-containing lysine demethylase associated with the Polycomb Repressive Complex 2 [PRC2] (Sasai *et al.* 2007; Herz *et al.* 2012).

To investigate a potential link between Su(z)2, Jarid2 and animal activity, we utilized the UAS/GAL4 system to knockdown the activity of the two proteins (Perkins *et al.* 2015). As complete loss of both Su(z)2 and Jarid2 is lethal (Kalisch and Rasmuson 1974; Sasai *et al.* 2007; Shalaby *et al.* 2017), we chose to explore knockdown in two tissues: Muscle, achieved by the *Mef2.R-GAL4* driver, which is expressed in somatic, visceral and cardiac muscle (Ranganayakulu *et al.* 1998); and neuronal tissues, achieved by the *elav^C155^-GAL4* driver, which is expressed in neurons starting at embryonic stage 12 (Lin and Goodman 1994). Because the knockdown with the *elav^C155^-GAL4* driver showed more impact in our hands, we focus on these data here. When animals lacking *Su(z)2* transcript in neurons are compared to their parents, either carrying the UAS-driven *Su(z)2* RNAi construct or the *elav^C155^-GAL4* driver, the animals with reduced *Su(z)2* transcript levels show significantly increased basal activity levels (Wilcoxon rank sum test, p=0.002797 for F1 compared to UAS parent, p=6.274e-06 for GAL4 parent). They also show significantly increased exercise-induced activity (Wilcoxon rank sum test, p=2.601e-07 for F1 compared to UAS parent, p=1.697e-05 for GAL4 parent), and the increase in activity is consistent in both for males and females (Figure 7). For *Jarid2*, we observe a similar increase in activity: if Jarid2 is removed in neuronal tissues, the animals show increased basal activity compared to their parents (UAS and GAL4 lines; Wilcoxon rank sum test, p=0.03412 for F1 compared to GAL4 parent, p=3.72e-06 for F1 compared to UAS parent). Exercised-induced activity is increased as well (Wilcoxon rank sum test, p= 0.02587 for F1 compared to GAL4 parent, p= 0.03752 for F1 compared to UAS parent). Together, these results indicate that our GWASs correctly identified *Su(z)2* and *Jarid2* as contributing to animal activity levels, both basal and exercise-induced.

**Figure 7.**
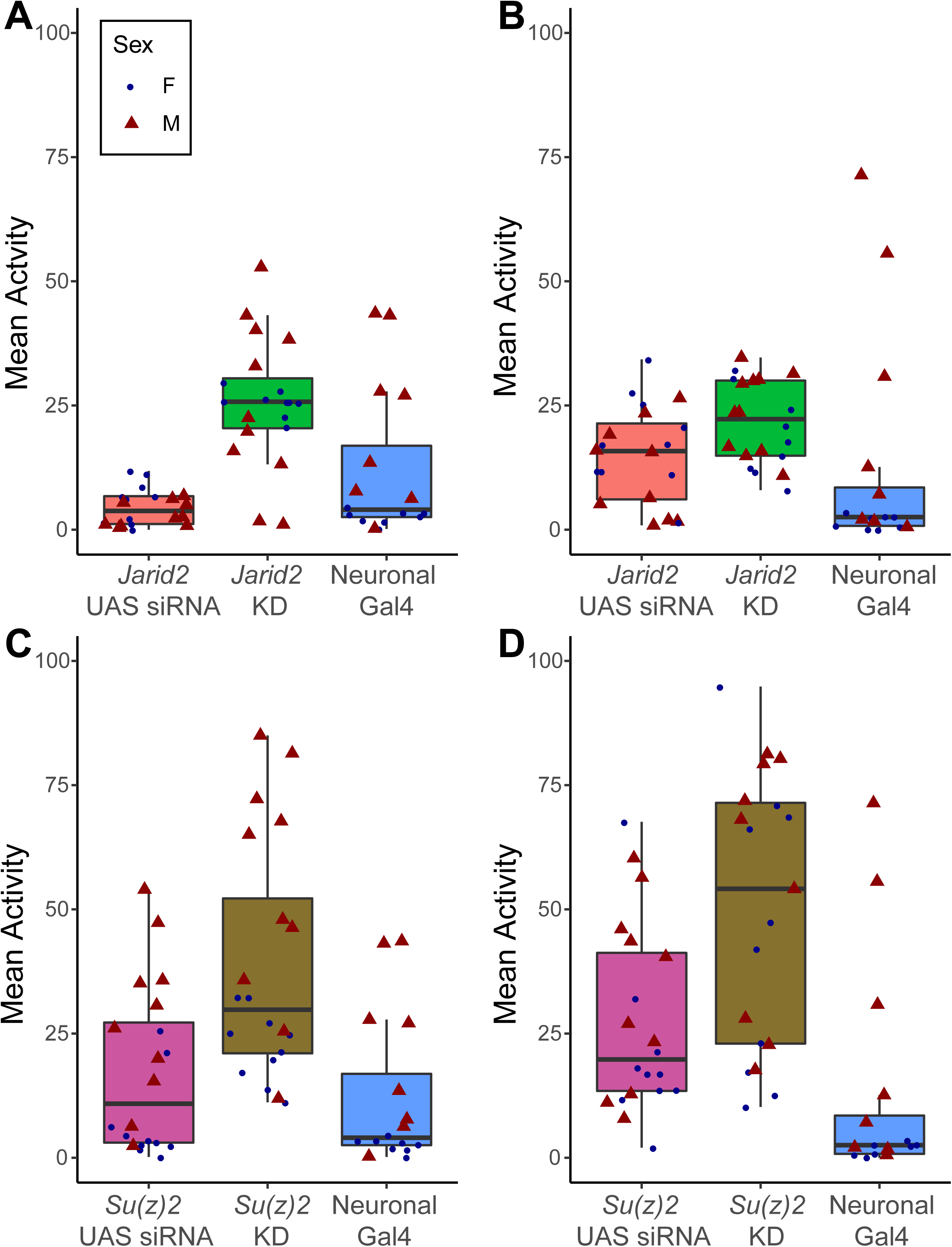
Knockdown of the polycomb group genes *Jarid2* and *Su(z)2* leads to increased activity levels. *Jarid2* and *Su(z)2* levels are decreased by expressing a UAS-controlled short hairpin construct targeting the gene of interest with a neuronal Gal4 driver (*elav*). In all boxplots, mean activity levels are plotted on the Y-axis, with data from males shown by a red triangle and data from females shown by a blue dot. **A.** Box plot showing the basal activity levels of animals with lower levels of *Jarid2* in their neuronal tissues (*Jarid2* KD, green) as well as their parents (UAS RNAi line, orange; elav-Gal4 line, blue). **B.** Box plot showing the induced activity levels of animals with lower levels of *Jarid2* in their neuronal tissues (*Jarid2* KD, green) as well as their parents (UAS RNAi line, orange; elav-Gal4 line, blue). **C.** Box plot showing the basal activity levels of animals with lower levels of *Su(z)2* in their neuronal tissues (*Su(z)2* KD, brown) as well as their parents (UAS RNAi line, purple; elav-Gal4 line, blue). **D.** Box plot showing the induced activity levels of animals with lower levels of *Su(z)2* in their neuronal tissues (*Su(z)2* KD, brown) as well as their parents (UAS RNAi line, purple; elav-Gal4 line, blue).

## DISCUSSION

As exercise is widely recommended as part of a healthy lifestyle and as a treatment for obesity, we explored the contribution of genetic variation to exercise in Drosophila using the DGRP2 strain collection. We found extensive variation in activity levels in this population, both basal and exercise-induced, which was dependent on genotype and sex. The GWASs identified over 300 genetic variants and more than 150 genes that contributed to the basal and exercise-induced activity phenotypes. Some of these genes had previously been associated with exercise performance or activity. This group of genes includes *couch potato* [*cpo*], an RNA-binding, nuclear protein expressed in the central nervous system that is essential for normal flight behavior (Bellen *et al.* 1992; Glasscock and Tanouye 2005; Schmidt *et al.* 2008). *nervous wreck* [*nwk*] encodes a very different kind of protein than *cpo*, but it also has been shown previously to impact animal activity: mutants in this FCH and SH3 domain-containing adaptor protein show paralysis due to problems at the synapse of neuromuscular junctions (Coyle *et al.* 2004; Rodal *et al.* 2008). Another gene in this group is *bedraggled* [*bdg*], which encodes a putative neurotransmitter transporter, and null mutants of which are described as flightless and uncoordinated (Rawls *et al.* 2007). The importance of the factors identified in our study is further underscored by the GO terms enrichment analysis, which revealed terms associated with the functions of the central nervous system and its interaction with the musculature. Together, these findings illustrate that the GWAS described here was successful in detecting variants associated with factors involved in the control of animal activity levels and behavior.

In addition to the genes such as *cpo*, *nwk*, and *bdg*, that would have been expected to impact basal activity or geotaxis-induced exercise activity, others candidate genes represent pathways with no clear link to animal activity or behavior. For example, variants in several chromatin proteins were identified as impacting animal activity levels, including SmydA-9, a SET domain protein, Su(z)2, a Polycomb group protein related to PSC, Jarid2, a Jumonji domain protein that interacts with PRC2, and Wde, an essential co-factor of the H3K9 methyltransferase Egg. In addition, several sperm proteins were identified as linked to activity (e. g. S-Lap8n, Sfp36F, Sfp51E) as were several members of the immunoglobulin superfamily (e. g. Side-II, CG31814, CG13506), but how they might contribute to an activity phenotype is unclear. As several of these genes are among a total of 36 genes identified here that were identified also by Schnorrer and colleagues as essential for normal muscle development (Schnorrer *et al.* 2010), it is likely they represent novel pathways linked to activity. Because it includes both unexpected and expected gene classes, the gene set identified here as contributing to both basal and exercise-induced activity levels provides a rich resource for researchers interested in using Drosophila as a tool for the study of exercise.

Our study also revealed that there were significant differences in the genes contributing to activity levels in males and in females, and that several genes could be identified that were responsible for the difference between the sexes. This finding was surprising, as we anticipated that the basic metabolic and sensory pathways involved in the control of animal activity would be conserved between males and females. Mackay and colleagues in 2012 discovered significant differences in the gene networks controlling starvation response and chill coma recovery time, but not startle response, and for all three traits, a large portion of the genetic variants that are identified in one sex show no significant association with the trait in the other sex (Ober *et al.* 2012). Garlapow and colleagues also found strong sex-specific differences in the factors controlling food intake in the DGRP2 population (Garlapow *et al.* 2015), and Morozova and colleagues detected sex differences in the networks controlling alcohol sensitivity (Fochler *et al.* 2017). These findings suggest that for many “basic” traits such as animal activity, sex specific differences in the underlying genetic networks occur frequently, highlighting the importance of studying both sexes.

When the list of candidate genes identified here is compared with those uncovered in other studies of activity traits, substantial overlap can be seen. For example, Jordan and colleagues investigated the genetic basis of the startle response and negative geotaxis in the DGRP2 population using a GWAS (Jordan *et al.* 2012). They identified approximately 200 genes associated with each of these activity phenotypes, a number of genes similar to that identified in our study. As our exercise system relies on negative geotaxis, not surprisingly, 24 genes identified by Jordan and colleagues were identified also in the basal activity analysis presented here, and an additional 17 genes from the exercise induced activity analysis overlapped with the gene set identified by Jordan and colleagues. Five genes were identified as candidates in all three analyses (*CG15630*, *CG33144*, *ed*, *nmo*, and *SKIP*) (Jordan *et al.* 2012). The overlap seen between the candidate genes identified in the two studies indicates that despite the different activity traits measured shared pathways exist.

Interestingly, a second activity study utilizing the DGRP2 identified a completely independent set of candidate genes, showing no overlap with the genes identified here. The study by Rhode and colleagues used video-tracking to monitor the activity of male Drosophila from the DGRP2 in a shallow petri dish by measuring distance traveled in a 5-minute interval (Rohde *et al.* 2018). This “2D” activity study focused on specific groups of genes linked to their phenotype, specifically genes involved in transmembrane transport. While this study used a very different algorithm to identify candidate genes, even when their phenotypic measures are analyzed with the standard GWAS tool our study used, we find no overlap in the candidate gene sets identified. This lack of overlap in candidate genes identified by two activity studies illustrates the complexity in the pathways that impact basic animal behaviors such as activity. Given current understanding, genes from basic energy metabolism pathways to genes controlling the development of muscles and sensory organs to genes impacting the processing of sensory information are all involved in controlling activity levels. Thus, it is not surprising that studies using different activity types will identify distinct sets of candidate genes; rather, these findings highlight that additional innovative studies are needed to come to a comprehensive understanding of the genes involved in animal activity, both basal and in response to stimulation.

The analysis of candidate genes presented here demonstrates that the REQS can be used successfully to identify genes involved in controlling basal and exercise induced activity levels in the DGRP2. Many of the candidate genes identified show relatively small impacts in the DGRP2, likely due to the presence of weak, not null alleles in this wild-derived population. This likelihood is especially high for genes where the null alleles are described as flightless, as such mutants are unlikely to survive outside the laboratory. Interestingly, while most of the candidate genes identified are sex-specific, sex dependencies are typically not described for flightless or flight defective alleles listed on Flybase. This observation suggests that sex differences in activity patterns and responses to stimuli might be negligible for strong or null alleles, but do become relevant for alleles with more subtle impacts and thus affect the ability to detect significant associations with a phenotype in GWASs.

The assays using *Jarid2* and *Suz(2)* knockdown suggest that these chromatin modifiers might indeed play a role in controlling exercise activity levels, and possibly exercise response. While to date, neither *Jarid2* nor *Su(z)2* have been linked directly to animal activity, there are additional data that support this finding. Several alleles of *Su(z)2* were reported that result in climbing defects or even climbing inability due to malformation of the adhesive pads on the legs of the animals (Husken *et al.* 2015). Thus, it is possible that other natural alleles exist that modify the ability of the animals to climb slippery surfaces such as those encountered in laboratory culture and assay vials. In addition, the transcription factor Mef2, which is responsible for normal muscle development (Taylor and Hughes 2017), appears to be impacted by changes in Jarid2: it is downregulated significantly in *Jarid2* mutant larvae [data from (Herz *et al.* 2012)], analyzed with GEO2R], suggesting that alterations in Jarid2 levels might lead to changes in muscle. Mef2 is also among the Jarid2 bound targets reported by Herz and colleagues, which generally are enriched significantly for GO terms related to the central nervous system [PANTHER analysis of data from (Herz *et al.* 2012)]. In addition, publicly available data on Flybase (Gramates *et al.* 2017) show that *Mef2* is enriched for H3K27 methylation (H3K27me3) in cells derived from mesoderm of 6-8hr old embryos (Bonn *et al.* 2012), suggesting that it might be under control of the polycomb system (as do several other studies (Marino and Di Foggia 2016)), providing another link between muscle development and the polycomb group proteins Jarid2 and Su(z)2. The polycomb system is reported to also play a role in the developing nervous system (Lomvardas and Maniatis 2016; Moccia and Martin 2018), which might be another explanation for the link between *Jarid2*, *Su(z)2*, and activity observed in this study. Thus, the literature suggests several mechanisms by which *Su(z)2* and *Jarid2* might impact animal activity due to their roles in the polycomb system of gene regulation.

In addition, changes in an organism’s activity levels have profound impacts, both acute and long-term, ranging from metabolic changes to physical changes to include psychological impacts in humans. Gene expression changes and alterations to the epigenome have been identified as immediate consequences of exercise (Hargreaves 2015; Soci *et al.* 2017). For example, there are several studies documenting DNA methylation changes as well as changes in histone acetylation following exercise in multiple systems (Voisin *et al.* 2015; Mcgee and Walder 2017). These epigenetic changes are one possible mechanism that might mediate the long-term consequences of exercise, many of which can persist even in the absence of further exercise. Thus, it is of interest that two proteins linked to the histone methylation mark H3K27me3 and the polycomb system were identified as contributing to the variation in activity levels between individuals in our study, and future studies in the role of these marks with regard to exercise are needed.

In summary, our study has identified promising candidate genes contributing to the variation in activity levels seen between genetically distinct individuals. Many of the genes identified have not been linked to exercise previously, and they thus present novel avenues for exploration to exercise biologists. Given the clear homology relationships between many of the Drosophila genes identified and mammalian genomes, the gene set presented here also provides new research directions for exercise biology studies in rodents. Using the results from model systems such as Drosophila as a guide, translational scientists will be able to accelerate biomarker development to eventually allow medical professionals to prescribe individualized exercise treatments for obesity-related diseases and to guide athletes of all kinds.

## ACKNOWLEDGEMENTS

The work described was supported partially by Award Number P30DK056336 from the National Institute Of Diabetes And Digestive And Kidney Diseases through a pilot grant from the UAB Nutrition and Obesity Research Center (NORC) to NCR. The content is solely the responsibility of the authors and does not necessarily represent the official views of the National Institute Of Diabetes And Digestive And Kidney Diseases or the National Institutes of Health.

We would like to thank our undergraduate students Michael Azar, Makayla Nixon, Jamie Hill, and Anthony Foster for their contributions to various aspects of these experiments. Dr. L. Reed (University of Alabama) graciously shared many of the DGRP2 strains with us. Additional stocks obtained from the Bloomington Drosophila Stock Center (NIH P40OD018537) were used in this study.

---

**Figure S1. Histograms illustrating the variability in activity levels among the lines of the DGRP2 population.** All lines binned based on their mean activity levels (average number of beam crossings recorded by the activity monitor in a 5-minute interval; bin size: 10). The bins are indicated on the X-axis (average number of beam crosses recorded by the activity monitor in a 5 minute interval), and the number of lines in each bin is shown on the Y-axis.

**A.** Histogram for basal activity in females.

**B.** Histogram for induced activity in females.

**C.** Histogram for basal activity in males.

**D.** Histogram for induced activity in males.

**Figure S2. Correlation between activity levels and lifespan.** To investigate the relationship between animal activity and lifespan, animal activity levels (X-axis, average number of beam crossings recorded by the activity monitor in a 5-minute interval) are plotted against life span (Y-axis, days) for each DGRP2 strain.

**A.** Basal activity.

**B.** Induced activity.

**Figure S3. Genetic variants associated with basal and induced activity levels in a GWAS using combined data from both sexes.** Chromosomal location of genetic variants (X-axis) are plotted against the negative log of the p-value testing for the likelihood of the variant being associated with the measured phenotype (Y-axis). The blue line in each plot marks the p=10^-5^ significance level, while the red line marks p=10^-7^.

**A.** Manhattan plot for basal activity.

**B.** Manhattan plot for induced activity.

**Figure S4. Small areas of linkage disequilibrium (LD) exist for the genetic variants associated with basal and exercise-induced activity.**

Heat maps illustrating the extent of LD among the genetic variants identified as significantly contributing to the basal (**A**) and exercise-induced (**B**) activity levels. The level of LD is indicated by the color of each square with red indicating the highest level of LD and blue indicating the absence of LD. The genetic variants are plotted based on their chromosomal location (X-and Y-axis), with the diagonal showing areas of with the strongest LD due to close physical proximity.

**Supplemental Table S1. Strains used for candidate gene analysis.** This table contains the genotypes and Bloomington stock numbers of the strains analyzed.

**Supplemental Table S2. List of DGRP lines included in this study.** Table is separated by treatment and sex for each line. Y = Yes, N = No.

**Supplemental Table S3. Phenotypic measures used in the GWAS analysis.** Summary data as submitted to the DGRP2 GWAS webtool as well as raw data are provided.

**Supplemental Table S4. Candidate genetic variants identified as significant by the GWAS.** This table includes results from the basal activity GWAS and the exercise-induced activity GWAS.

**Supplemental Table S5. Detailed GO term enrichment analysis results for genetic variants associated with basal activity.** This table includes the complete GO term enrichment analysis results for the “Cellular component” and “Biological Process” categories for genetic variants associated with basal activity.

