## Supplemental Table S2 for "Genetic networks underlying natural variation in basal and induced activity levels in *Drosophila melanogaster*"

**Supplemental Table 2. DGRP2 lines included in this study.**

| **Genotype** | **M** | **F** | **Basal** | **Induced** |
| --- | --- | --- | --- | --- |
| 21 | Y | Y | Y | Y |
| 26 | Y | Y | Y | Y |
| 28 | Y | Y | Y | Y |
| 31 | Y | Y | Y | Y |
| 38 | Y | Y | N | Y |
| 40 | Y | Y | Y | Y |
| 41 | Y | Y | Y | Y |
| 42 | Y | Y | Y | Y |
| 45 | Y | Y | Y | Y |
| 48 | Y | Y | N | Y |
| 49 | Y | Y | Y | Y |
| 57 | Y | Y | Y | N |
| 59 | Y | Y | Y | Y |
| 69 | Y | Y | Y | Y |
| 73 | Y | Y | Y | Y |
| 83 | Y | Y | Y | Y |
| 85 | Y | Y | Y | Y |
| 88 | Y | Y | Y | Y |
| 93 | Y | Y | N | Y |
| 100 | Y | Y | Y | Y |
| 101 | Y | Y | Y | Y |
| 105 | Y | Y | Y | Y |
| 109 | Y | Y | Y | N |
| 129 | Y | Y | Y | Y |
| 138 | Y | Y | Y | Y |
| 142 | Y | Y | Y | Y |
| 149 | Y | Y | Y | Y |
| 153 | Y | Y | Y | Y |
| 158 | Y | Y | Y | Y |
| 161 | Y | Y | Y | N |
| 177 | Y | Y | Y | Y |
| 181 | Y | Y | Y | Y |
| 195 | Y | Y | N | Y |
| 227 | Y | Y | Y | Y |
| 228 | Y | Y | Y | Y |
| 229 | Y | Y | Y | Y |
| 233 | Y | Y | Y | Y |
| 235 | Y | Y | Y | N |
| 256 | Y | Y | N | Y |
| 287 | Y | Y | Y | Y |
| 301 | Y | Y | Y | Y |
| 304 | Y | Y | Y | Y |
| 306 | Y | Y | Y | Y |
| 307 | Y | Y | Y | Y |
| 309 | Y | Y | Y | Y |
| 310 | Y | Y | Y | Y |
| 313 | Y | Y | Y | Y |
| 315 | Y | Y | Y | Y |
| 317 | Y | Y | Y | Y |
| 318 | Y | Y | Y | Y |
| 320 | Y | Y | Y | Y |
| 321 | Y | Y | Y | Y |
| 324 | Y | Y | Y | Y |
| 332 | Y | Y | Y | Y |
| 335 | Y | Y | Y | Y |
| 336 | Y | Y | Y | Y |
| 338 | Y | Y | Y | Y |
| 340 | Y | Y | Y | Y |
| 350 | Y | Y | Y | Y |
| 352 | Y | Y | Y | Y |
| 357 | Y | Y | Y | Y |
| 358 | Y | Y | Y | Y |
| 359 | Y | Y | Y | Y |
| 360 | Y | Y | Y | Y |
| 361 | Y | Y | Y | Y |
| 362 | Y | Y | Y | Y |
| 365 | Y | Y | Y | Y |
| 367 | Y | Y | N | N |
| 370 | Y | Y | Y | Y |
| 371 | Y | Y | Y | Y |
| 373 | Y | Y | Y | Y |
| 374 | Y | Y | Y | Y |
| 375 | Y | Y | Y | Y |
| 377 | Y | Y | Y | Y |
| 379 | Y | Y | Y | Y |
| 380 | Y | Y | Y | N |
| 381 | Y | Y | Y | Y |
| 382 | Y | Y | Y | Y |
| 383 | Y | Y | Y | Y |
| 386 | Y | Y | N | Y |
| 390 | Y | Y | Y | Y |
| 391 | Y | Y | Y | Y |
| 395 | Y | Y | Y | Y |
| 397 | Y | Y | Y | Y |
| 399 | Y | Y | Y | Y |
| 405 | Y | Y | Y | Y |
| 406 | Y | Y | Y | Y |
| 409 | Y | Y | Y | Y |
| 426 | Y | Y | Y | Y |
| 427 | Y | Y | Y | Y |
| 439 | Y | Y | Y | N |
| 441 | Y | Y | Y | N |
| 443 | Y | Y | Y | Y |
| 461 | Y | Y | Y | Y |
| 486 | Y | Y | Y | Y |
| 492 | Y | Y | Y | Y |
| 502 | Y | Y | Y | Y |
| 505 | Y | Y | Y | Y |
| 508 | Y | Y | Y | Y |
| 509 | Y | Y | Y | Y |
| 513 | Y | Y | Y | Y |
| 528 | Y | Y | Y | Y |
| 530 | Y | Y | Y | Y |
| 535 | Y | Y | Y | Y |
| 555 | Y | Y | Y | Y |
| 559 | Y | Y | Y | Y |
| 589 | Y | Y | Y | Y |
| 595 | Y | Y | Y | Y |
| 596 | Y | Y | Y | Y |
| 627 | N | Y | N | N |
| 634 | Y | Y | Y | Y |
| 639 | Y | Y | Y | Y |
| 703 | Y | Y | Y | Y |
| 705 | N | Y | N | N |
| 707 | Y | Y | Y | Y |
| 714 | Y | Y | Y | Y |
| 716 | Y | Y | Y | Y |
| 727 | N | Y | N | Y |
| 737 | Y | Y | Y | Y |
| 738 | Y | Y | Y | Y |
| 748 | Y | Y | N | Y |
| 757 | Y | Y | Y | Y |
| 761 | Y | Y | Y | Y |
| 765 | Y | Y | Y | Y |
| 774 | Y | Y | Y | Y |
| 783 | Y | Y | Y | Y |
| 786 | Y | Y | Y | Y |
| 787 | Y | Y | Y | Y |
| 790 | Y | Y | Y | Y |
| 796 | Y | Y | Y | Y |
| 799 | Y | Y | Y | Y |
| 801 | Y | Y | Y | Y |
| 805 | Y | Y | Y | Y |
| 808 | Y | Y | Y | Y |
| 810 | Y | Y | Y | Y |
| 812 | Y | Y | Y | Y |
| 818 | Y | Y | Y | N |
| 819 | Y | Y | Y | Y |
| 820 | Y | Y | Y | Y |
| 821 | Y | Y | Y | Y |
| 822 | Y | Y | Y | Y |
| 832 | Y | Y | Y | Y |
| 837 | N | Y | Y | Y |
| 843 | Y | Y | Y | Y |
| 849 | Y | Y | Y | Y |
| 850 | Y | Y | Y | N |
| 852 | Y | Y | Y | Y |
| 853 | Y | Y | Y | Y |
| 855 | Y | Y | Y | Y |
| 857 | Y | Y | Y | Y |
| 859 | Y | Y | Y | Y |
| 879 |  |  | Y | Y |
| 882 | Y | Y | Y | Y |
| 887 | Y | Y | Y | Y |
| 890 | Y | Y | Y | Y |
| 892 | Y | Y | Y | Y |
| 897 | Y | Y | Y | Y |
| 900 | Y | Y | Y | Y |
| 908 | Y | Y | Y | Y |
| 911 | Y | Y | Y | N |
| 913 | Y | Y | Y | Y |
