## Supplementary figures and images for "Genetic networks underlying natural variation in basal and induced activity levels in *Drosophila melanogaster*"

### Supplemental Figure S1

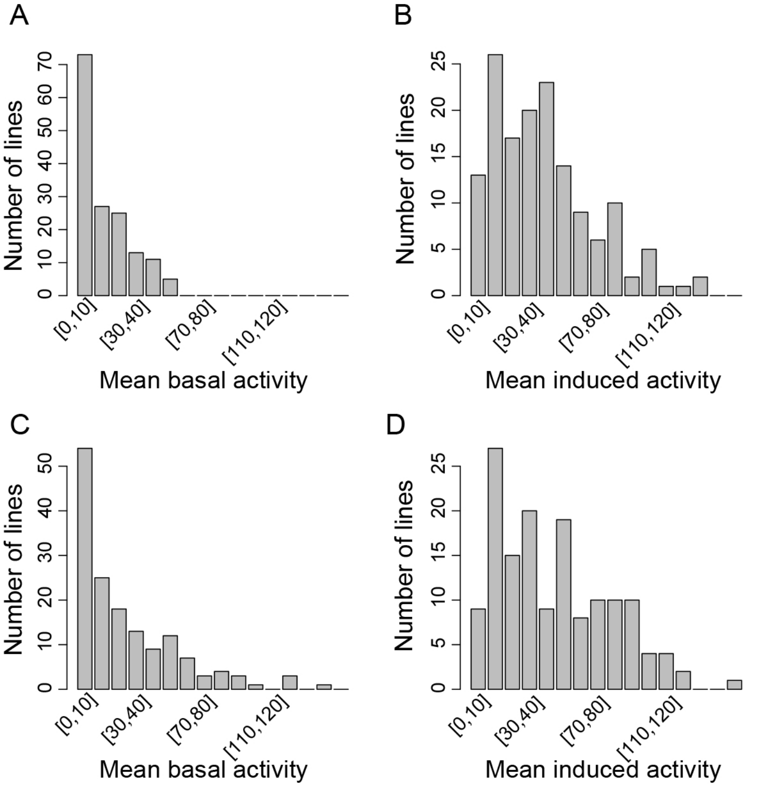

### Supplemental Figure S2

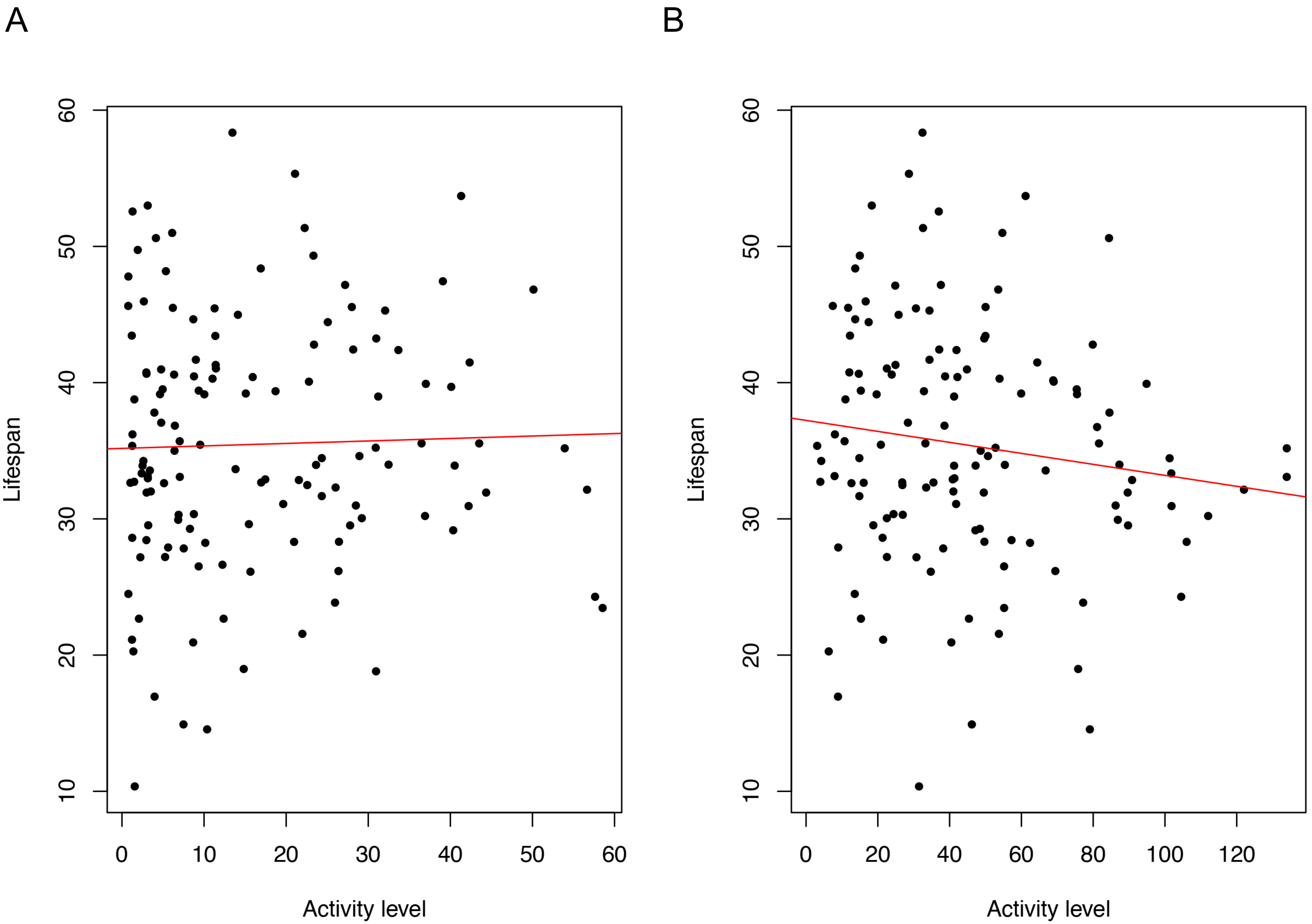

### Supplemental Figure S3

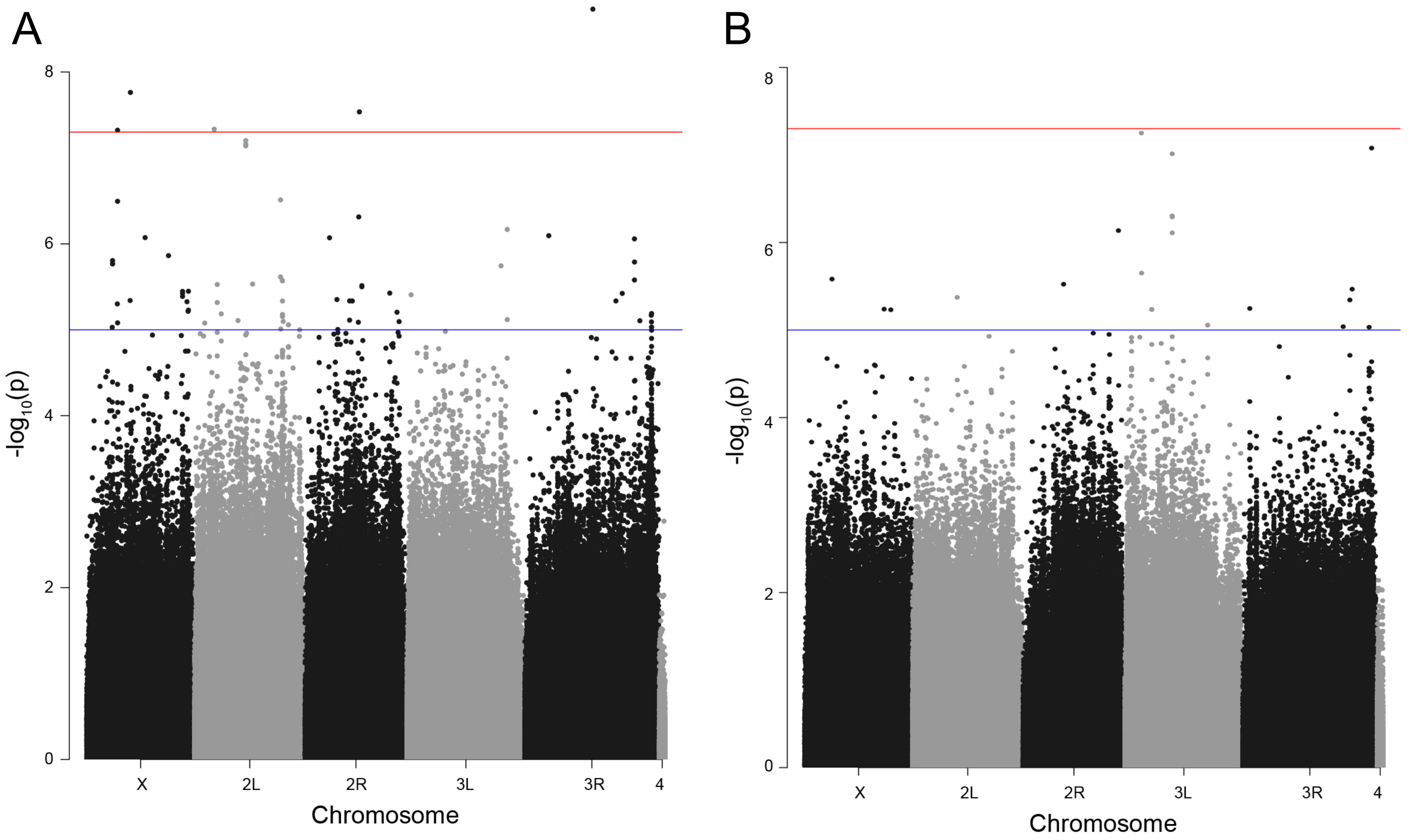

### Supplemental Figure S4

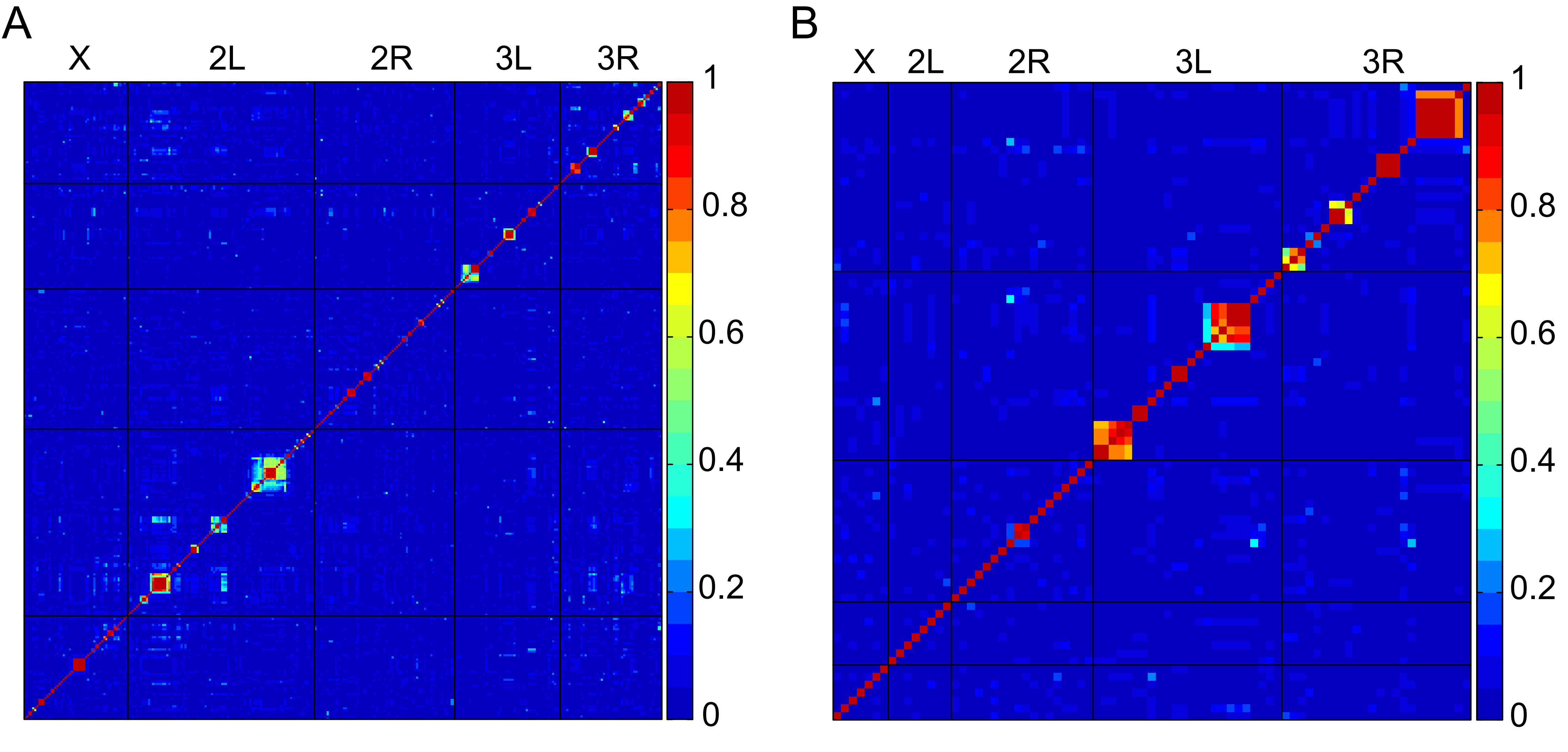
